# Social information-mediated population dynamics in non-grouping prey

**DOI:** 10.1101/2022.03.21.484882

**Authors:** Zoltán Tóth, Gabriella Csöppü

## Abstract

Inadvertent social information (ISI) use, i.e., the exploitation of social cues including the presence and behaviour of others, has been predicted to mediate population-level processes even in the absence of cohesive grouping. However, we know little about how such effects may arise when the prey population lacks social structure beyond the spatiotemporal autocorrelation originating from the random movement of individuals. In this study, we built an individual-based model where predator avoidance behaviour could spread among randomly moving prey through the network of nearby observers. We qualitatively assessed how ISI use may affect prey population size when cue detection was associated with different probabilities and fitness costs, and characterised the structural properties of the emerging detection networks that would provide pathways for information spread in prey. We found that ISI use was among the most influential model parameters affecting prey abundance and increased equilibrium population sizes in most examined scenarios. Moreover, it could substantially contribute to population survival under high predation pressure, but this effect strongly depended on the level of predator detection ability. When prey exploited social cues in the presence of high predation risk, the observed detection networks consisted of a large number of connected components with small sizes and small ego networks; this resulted in efficient information spread among connected individuals in the detection networks. Our study provides hypothetical mechanisms about how temporary local densities may allow information diffusion about predation threats among conspecifics and facilitate population stability and persistence in non-grouping animals.

**Significance Statement:** The exploitation of inadvertently produced social cues may not only modify individual behaviour but also fundamentally influence population dynamics and species interactions. Using an individual-based model, we investigated how the detection and spread of adaptive antipredator behaviour may cascade to changes in the demographic performance of randomly moving (i.e., non-grouping) prey. We found that social information use contributed to population stability and persistence by reducing predation-related per capita mortality and raising equilibrium population sizes when predator detection ability reached a sufficient level. We also showed that temporary detection networks had structural properties that allowed efficient information spread among prey under high predation pressure. Our work represents a general modelling approach that could be adapted to specific predator-prey systems and scrutinize how temporary local densities allow dynamic information diffusion about predation threats and facilitate population stability in non-grouping animals.

## Introduction

Organisms have to gather information about their surroundings to overcome challenges such as finding resources and avoiding danger (Dall and Johnstone 2002). For that, individuals directly interact with the environment to gain up-to-date information about its state (‘personal information’), but they can also complement that knowledge by utilizing social information for optimal decision-making (Galef and Giraldeau 2001; Dall et al. 2005; Bonnie and Earley 2007; Hoppitt and Laland 2013). One type of social information is associated with inadvertently produced social cues that include the presence or the behaviour of others, or the product of their behaviour such as scent marks, excretions or food remnants, all of which may provide relevant information about current environmental conditions. Inadvertent social information (ISI) use is known to occur in many ecological contexts, including predator avoidance, foraging and habitat choice (Danchin et al. 2004; Gil et al. 2018). The advantages of living in social groups are thought to include the opportunity to access social information (Krause and Ruxton 2002; Ward and Webster 2016; Goodale et al. 2017), and thus ISI use is usually associated with species where social interactions promote information transmission among group-mates (King and Cowlishaw 2007; Duboscq et al. 2016; Gil et al. 2017).

Under predation risk, dynamic information about threats is transmitted from alarmed group members to naïve ones, a phenomenon that is commonly called collective detection (Lima 1990; Pays et al. 2013). This process often takes place through evolved signals such as alarm calls, but social cues including sudden movements (Coleman 2008; Hingee and Magrath 2009; Boujja-Miljour et al. 2017), fright responses (Chivers and Ferrari 2014; Cruz et al. 2020), or changes in posture (Brown et al. 1999; Pays et al. 2013) have also been found to convey information about the presence of predators in animal collectives. Adjustments to the behaviour of others (also referred to as ‘behavioural contagion’; Firth 2020) do not only affect individual fitness by increasing survival probabilities, but can also lead to the emergence of correlated behaviours and space use in many individuals and thus influence system-level functions (Goodale et al. 2010; Gil et al. 2018; Tóth 2021). Previous theoretical models have predicted that ISI use can prevent population collapses under high predation pressure (Gil et al. 2017, 2018) and facilitate the coexistence of competing species that share common predators (Parejo and Avilés 2016; Gil et al. 2019). Empirical evidence also indicates that the utilization of social information can influence the material flux on the ecosystem level (Gil and Hein 2017). By promoting adaptive behavioural responses to environmental uncertainties (e.g., due to anthropogenic effects [Greggor et al. 2017], in the distribution of resources [O’Mara et al. 2014] or predation risk [Crane et al. 2022]), ISI use has the potential to minimize the impact of morphological, physiological or genetic adaptations (Laland 1992) or influence genetic change through gene–culture coevolution (Whitehead et al. 2019).

There are animal species that do not exhibit social attraction toward conspecifics and therefore do not form permanent or periodical cohesive groups. We refer to these organisms as non-grouping animals (for more details about to this definition, see Tóth et al. 2020). Lacking motivation for social cohesion, non-grouping animals do not maintain spatial proximity with others, and thus direct interactions between conspecifics can be infrequent. Nevertheless, such individuals may also exploit social cues (e.g., visual, acoustic, chemical or vibrational cues) when these are within the range of relevant sensory perception. Moreover, social information may also diffuse among nearby observers via ‘detection networks’ (reviewed in Tóth et al. 2020). If so, spatial changes in social cues over time (e.g., relative differences in activity and associated conspicuousness; Chivers and Ferrari 2014) can provide dynamic information about predation threats in many terrestrial and aquatic systems (Gil et al. 2017). In accordance with this idea, wood crickets (*Nemobius sylvestris*) adaptively change their behaviour after having observed the predator avoidance behaviour of knowledgeable conspecifics, and this information is transmitted to and utilized by other naïve individuals as well (Coolen et al. 2005). In temporary aggregations, escape responses of Iberian green frogs (*Rana perezi*) are also influenced by the behaviour of adjacent conspecifics (Martín et al. 2006). In mixed-species aggregations of non-schooling fish, the density (number of fish in the foraging area) and behaviour (when to feed in and when to flee from the foraging area) of nearby individuals are being used as inadvertent social information (Gil and Hein 2017). The resulting behavioural coupling among individuals, in turn, affects both species abundance and the amount of algae consumed and as a result, determines the total material flow in the coral reef ecosystem. While such observations prove that threat-related social cues can be exploited by non-grouping animals in some instances, the general conditions under which ISI use exerts a positive effect on population stability and persistence in such species have remained largely unexplored. For example, thresholds associated with the cost of antipredator behaviour and probabilities of cue detection (i.e., the detection of predators or conspecifics’ behaviour) may set boundaries for social information-modulated population-level effects under different predation pressure regimes. Similarly, detection networks may have only a limited capacity to provide efficient information pathways for the emergence of such effects. In a previous work, Tóth (2021) used an individual-based model to test specifically how ISI use may alter the relationship between fluctuating predator and non-grouping prey populations. The author found that ISI use can disrupt population cycles and decrease the strength of second-order density dependence between predator and prey (i.e., negative feedback of the second order that would imply an interaction between predator and prey populations in either direction), thus stabilize their dynamics and facilitate their long-term coexistence.

In this study, we investigated how the detection and spread of predator avoidance behaviour among conspecifics affected demographic performance in non-grouping prey. We constructed an individual-based model of prey and generalist predator populations where individuals (both prey and predators) moved randomly on the landscape, and social information could diffuse through the observation of antipredator behaviour in prey. This model, an extension of our earlier model presented by Tóth (2021), allowed us to assess qualitatively how ISI use may cascade to population-level changes in noncyclic prey populations that experience relatively constant predation pressures and lack social structure. We could also examine the structural properties of detection networks that might facilitate the emergence of such effects. Thus, predictions from the presented model may be applied more generally to non-grouping organisms compared to those from our previous work.

## Materials and Methods

### Model construction

We simulated a homogeneous, continuous 2D landscape (80 × 80 spatial units) where both prey and predators moved randomly by exhibiting correlated random walks (CRW). CRW considers short-term correlations between successive step orientations and has been successfully used to model animals’ non-orientated search paths for a long time (Benhamou 2006; Codling et al. 2008; Reynolds 2014). In CRW models, habitat must be rather homogeneous; examples of such natural habitats include beaches and deserts, grasslands, agricultural crops, or those where resource patch distribution at the large scale is uniform or random (Byers 2001). At the start of a simulation cycle, 500 prey and 150 predators were randomly placed on the landscape, and then individuals performed a given set of behaviours (Fig. 1, Table 1). During movement, each individual’s movement distance was randomly selected between zero and a maximum value given by the parameters *d*_prey_ and *d*_P_ for prey and predators, respectively. Turning angles were determined by random deviates drawn from wrapped Cauchy circular distribution with *μ*=0 and *ρ*=0.8. At the landscape edge, individuals moved to the opposite side of the landscape when crossing a boundary and continued moving (i.e., torus landscape with no edge). Both prey and predator could also detect other individuals through the landscape edge. We assumed that only one individual could survive within the range of one spatial unit due to competition in both prey and predators (after movement and dispersion of offspring; see Fig. 1), introducing density-dependent mortality in their populations. In this system, we assumed non-dynamic predators that can exert high predation pressure on the prey population, thus predator population size was determined only by their reproductive rate and density-dependent mortality, but was unaffected by the success of hunting (as if switching to alternative prey when necessary). Consequently, predator and prey populations were noncyclic and demographically decoupled (for a similar approach, see Gil et al. 2019), and prey populations experienced predation pressures that were directly proportional to the given value of predators’ reproduction-related parameter (Table 1).

**Fig. 1.**
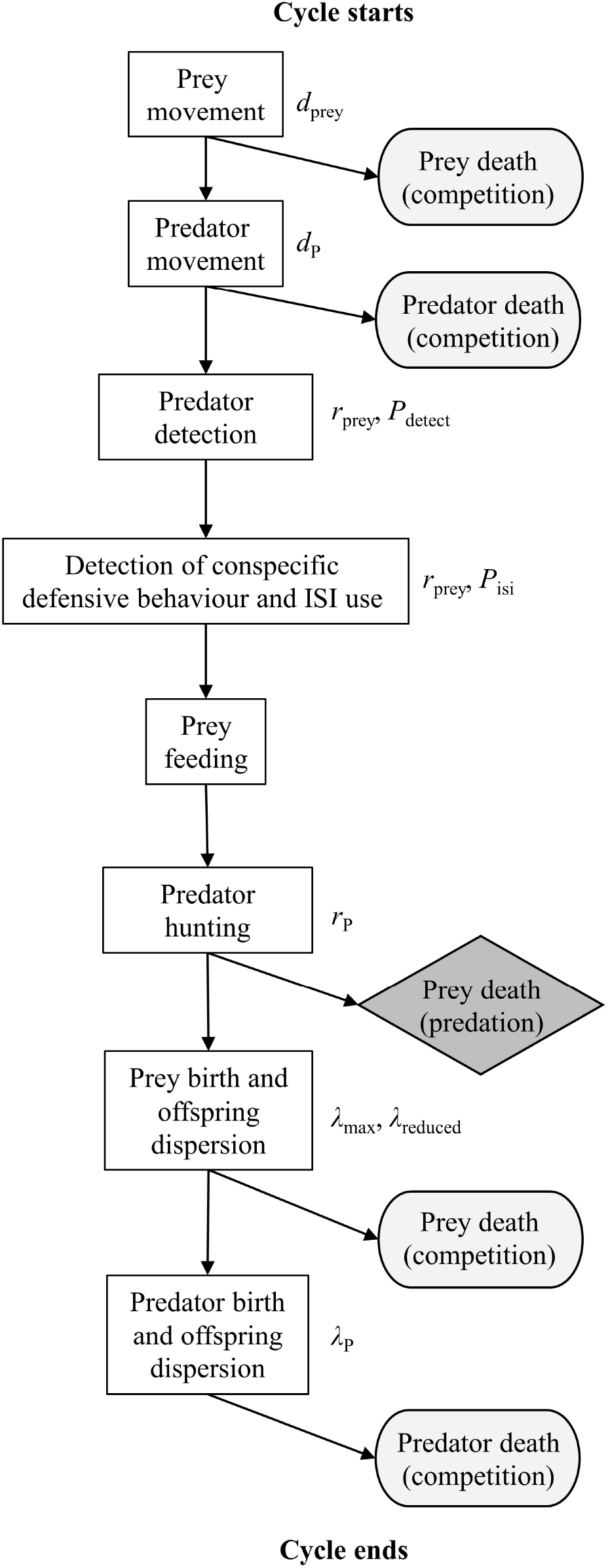
Model flowchart for a single simulation cycle. Sequential prey and predator behaviours are listed together with the model parameter(s) associated with the given steps. Behavioural steps resulting in a decrease in population size, i.e., mortality due to intraspecific competition (in rounded rectangles) or predation (in diamond) are shown in light and dark grey, respectively

**Table 1.**
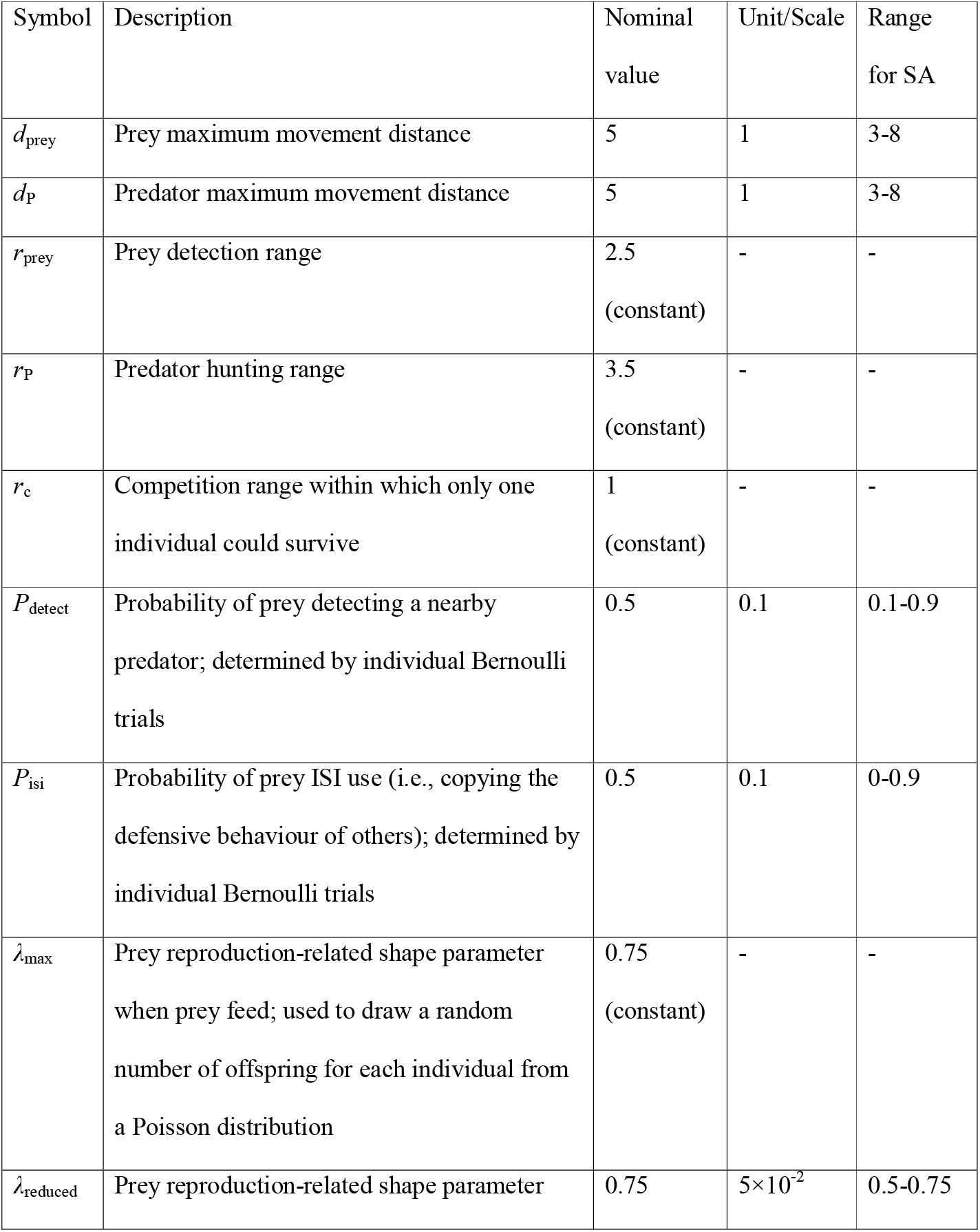
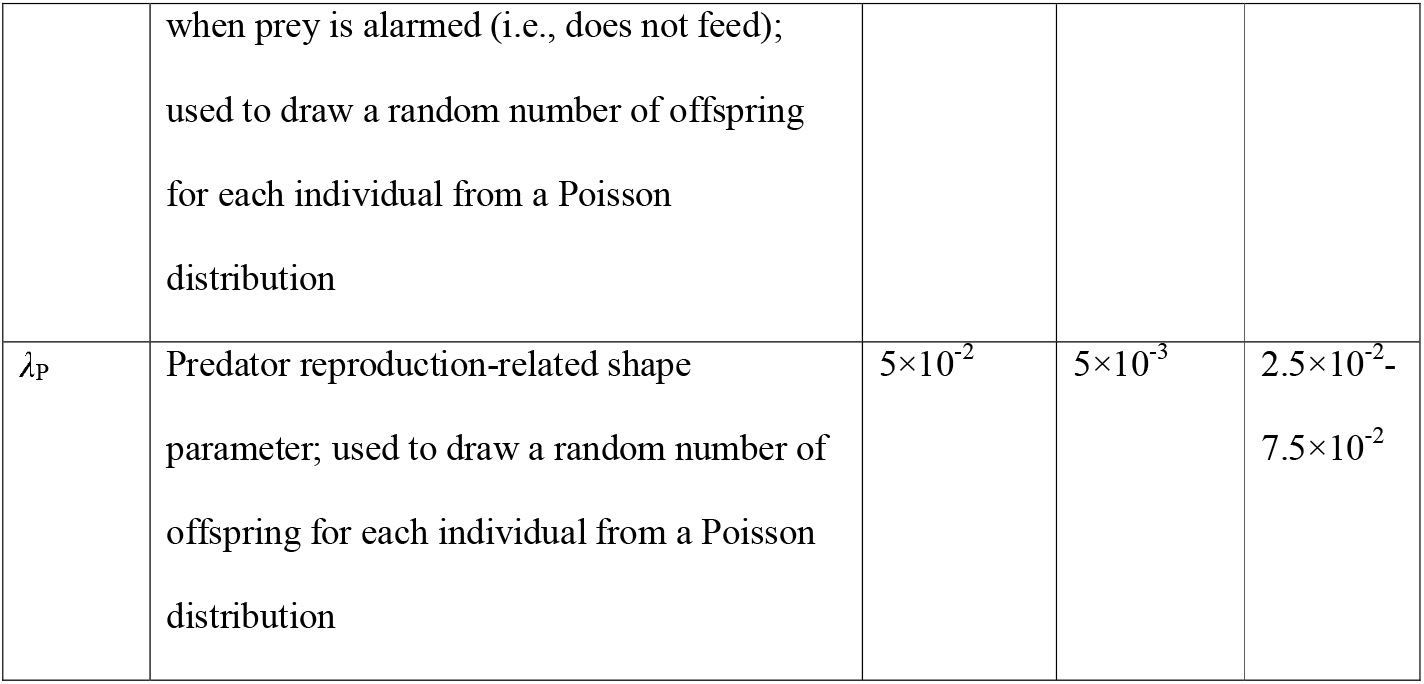
Model parameters and their range for sensitivity analysis (SA). Maximum movement distances indicate the maximum number of spatial units that an individual could travel on the landscape in a simulation cycle; the actual integer value was randomly selected between zero and this maximum value

| Symbol | Description | Nominal value | Unit/Scale | Range for SA |
| --- | --- | --- | --- | --- |
| $d_{\text{prey}}$ | Prey maximum movement distance | 5 | 1 | 3-8 |
| $d_{\text{p}}$ | Predator maximum movement distance | 5 | 1 | 3-8 |
| $r_{\text{prey}}$ | Prey detection range | 2.5<br>(constant) | - | - |
| $r_{\text{p}}$ | Predator hunting range | 3.5<br>(constant) | - | - |
| $r_{\text{c}}$ | Competition range within which only one individual could survive | 1<br>(constant) | - | - |
| $P_{\text{detect}}$ | Probability of prey detecting a nearby predator; determined by individual Bernoulli trials | 0.5 | 0.1 | 0.1-0.9 |
| $P_{\text{isi}}$ | Probability of prey ISI use (i.e., copying the defensive behaviour of others); determined by individual Bernoulli trials | 0.5 | 0.1 | 0-0.9 |
| $\lambda_{\text{max}}$ | Prey reproduction-related shape parameter when prey feed; used to draw a random number of offspring for each individual from a Poisson distribution | 0.75<br>(constant) | - | - |
| $\lambda_{\text{reduced}}$ | Prey reproduction-related shape parameter when prey is alarmed (i.e., does not feed);<br>used to draw a random number of offspring<br>for each individual from a Poisson<br>distribution | 0.75 | $5 \times 10^{-2}$ | 0.5-0.75 |
| $\lambda_p$ | Predator reproduction-related shape<br>parameter; used to draw a random number of<br>offspring for each individual from a Poisson<br>distribution | $5 \times 10^{-2}$ | $5 \times 10^{-3}$ | $2.5 \times 10^{-2}$-<br>$7.5 \times 10^{-2}$ |

In the absence of predators, prey moved, competed, fed and reproduced in the simulated landscape. Prey population size resulted in this scenario was regarded as being in equilibrium at the carrying capacity of the environment. When present, each predator could consume a maximum of five prey individuals in a cycle within its hunting range, which was defined as an *r_P_* distance from the predator’s position in any direction. Prey could detect predators that were *r*_prey_ distance with a probability given by *P*_detect_ (determined by individual Bernoulli trials). Thus, the detection of other individuals (either predators, prey or conspecifics) depended on individual sensory ranges that also crossed through the torus landscape edges, bringing more reality to the simulation compared to the model presented in Tóth (2021). Upon successfully detecting a predator within *r*_prey_ distance, prey became alarmed and hid, and thus were undetectable to predators. However, these individuals did not feed either and consequently could have a reduced reproduction rate. Thus, prey animals were capable of behaviourally adjusting their exposure to predators (with the probability ranging between 0.1 and 0.9; see Table 1), but this antipredator behaviour potentially incurred a fitness cost. Lima and Dill (1990) summarized supporting evidence for costly behavioural responses to predation risk in multiple taxa, including a reduction in feeding, growth or reproduction. Predators hunted on visible, feeding prey with a 50% success (determined by individual Bernoulli trials). Similarly high success rates against some prey species have been observed in generalist predators such as red foxes (*Vulpes vulpes*; Červený et al. 2012), *Drassodes lapidosus* spiders (Michálek et al. 2017), or American kestrels (*Falco sparverius;* Toland 1987). Prey could also detect predators indirectly by observing alarmed conspecifics within *r*_prey_ distance with a probability given by *P*_isi_ (determined by individual Bernoulli trials). Being alarmed had the same consequences (i.e., immune to predation, reduced reproduction rate) irrespective of the detection mode. Wide ranges of possible values for both *P*_detect_ and *P*_isi_ (see in Table 1) were used to cover most scenarios in which ISI use may occur under natural conditions. We did not manipulate cue reliability in the model, we simply considered that ISI use had a higher cost when social cues could also be false and individuals responded to those indiscriminately. Prey feeding occurred once in a cycle in prey that was not hiding. The number of offspring for each individual was sampled from a Poisson distribution with the shape parameter given by *λ*_reduced_ for alarmed prey, *λ*_max_ for fed prey, and *λ*_P_ for predators in each cycle. Offspring dispersed in the same cycle 8, 9 or 10 spatial units away (randomly chosen) from the parent in both prey and predators. These higher step values (10 spatial units is the double of maximum *d*_prey_ and *d*_P_) were chosen to reflect that juvenile dispersion distances can far exceed adult movement ranges.

### Detection networks

From the spatial distribution of prey, we defined detection networks based on the range within which individuals could observe the behaviour of others (i.e., exploit social cues if present) in each simulation cycle (Fig. 2). In such networks, nodes represent individuals, and edges denote the possibility of mutual observation. If a prey individual became alarmed because it successfully detected a predator, information could spread from this individual to other conspecifics in the network under the following rules. The probability of information acquisition from one node to another is given by *w^k^*, where *w* is the edge weight (corresponding to the probability of information spread from one node to another through the edge between them and specified by the parameter *P*_isi_ in the model) and *k* is the number of steps on the shortest path between the two nodes. Only shortest paths were used to minimize the “travel time” of information between nodes in the network. During simulations, the maximum number of steps between the focal and the observed nodes was set to two and the total number of observed neighbours to ten (i.e., *k*_max_=2 and ∑(*n*)_max_=5 in each *k* step). Thus, an individual could receive information from a maximum of ten of its neighbours that were a maximum of two steps away in the detection network. With such restrictions, ISI use did not facilitate the emergence of large aggregations in prey and did not occur far outside the hunting range of predators. If there were more than five nodes at *k* step to a focal node, we randomly selected five. For any individual, the total probability of receiving information from its neighbours was calculated using the inclusion-exclusion principle (Allenby and Slomson 2010).

**Fig. 2.**
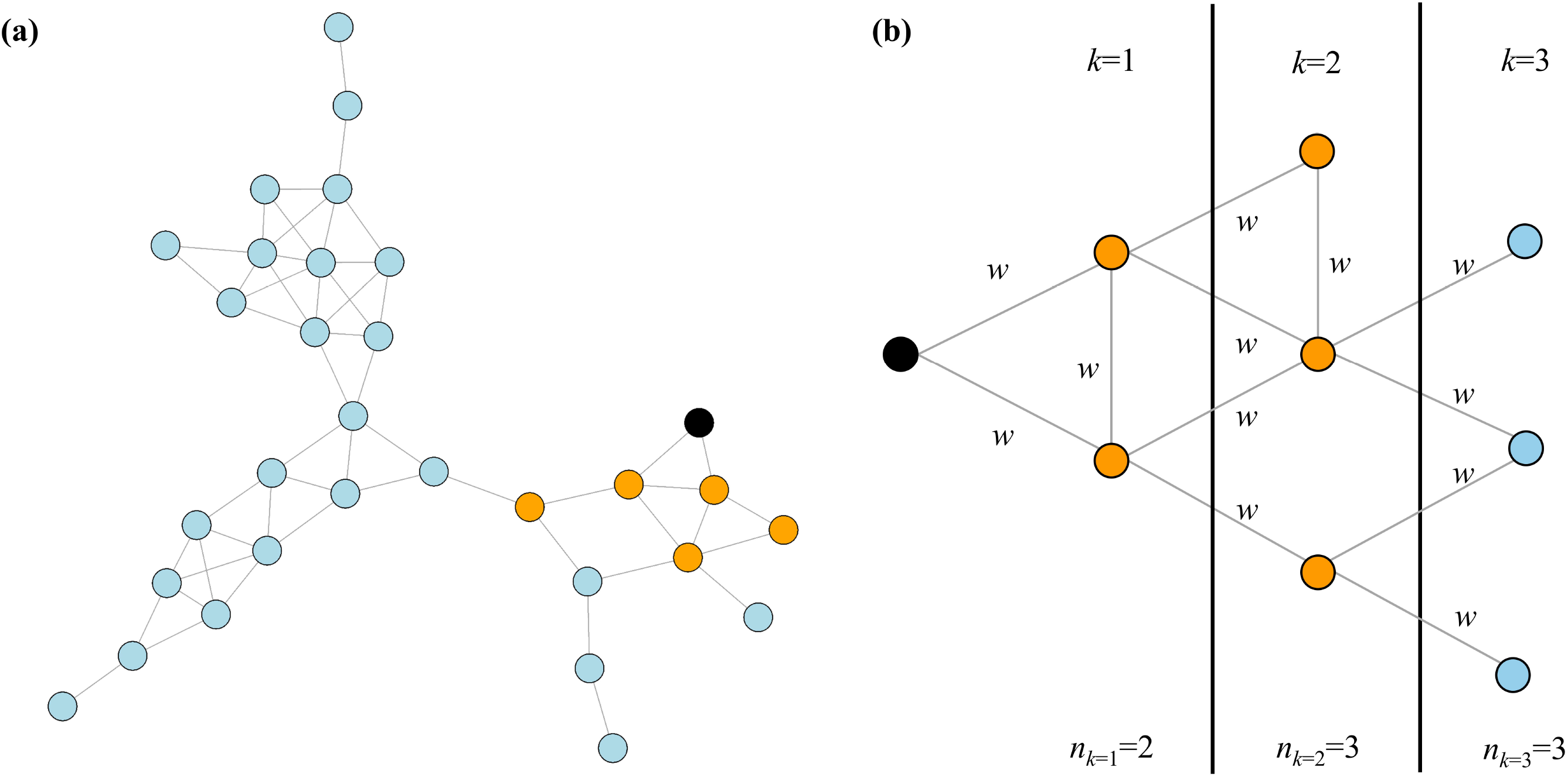
Schematic figure of a detection network (a) and segment of an individual ego network embedded within that network (b). Nodes represent individuals and edges denote the possibility of mutual observation. The probability of information acquisition from one node to another is given by *w^k^*, where *w* is the edge weight and *k* is the number of steps on the shortest path between the two nodes. For any individual, the total probability of receiving information from neighbours is calculated using the inclusion-exclusion principle. In our model, we used the settings *k*_max_=2 and ∑*n*_max_=5 in each *k* step, so the focal individual (black circle) could receive social information from a maximum of ten neighbours that were a maximum of two steps away in the detection network (orange circles)

### Analysis of simulation outputs

All simulations and calculations were performed in R 4.0.4 (R Core Team 2021). Instead of frequentist hypothesis testing, we focused on evaluating the magnitude of differences between simulation runs with different parameter settings (White et al. 2014). We ran the population simulations for 200 cycles (this interval was sufficient to reach equilibrium prey population size in the studied scenarios; see in Fig. 3a) and used the data from the last cycle in all calculations.

**Fig. 3.**
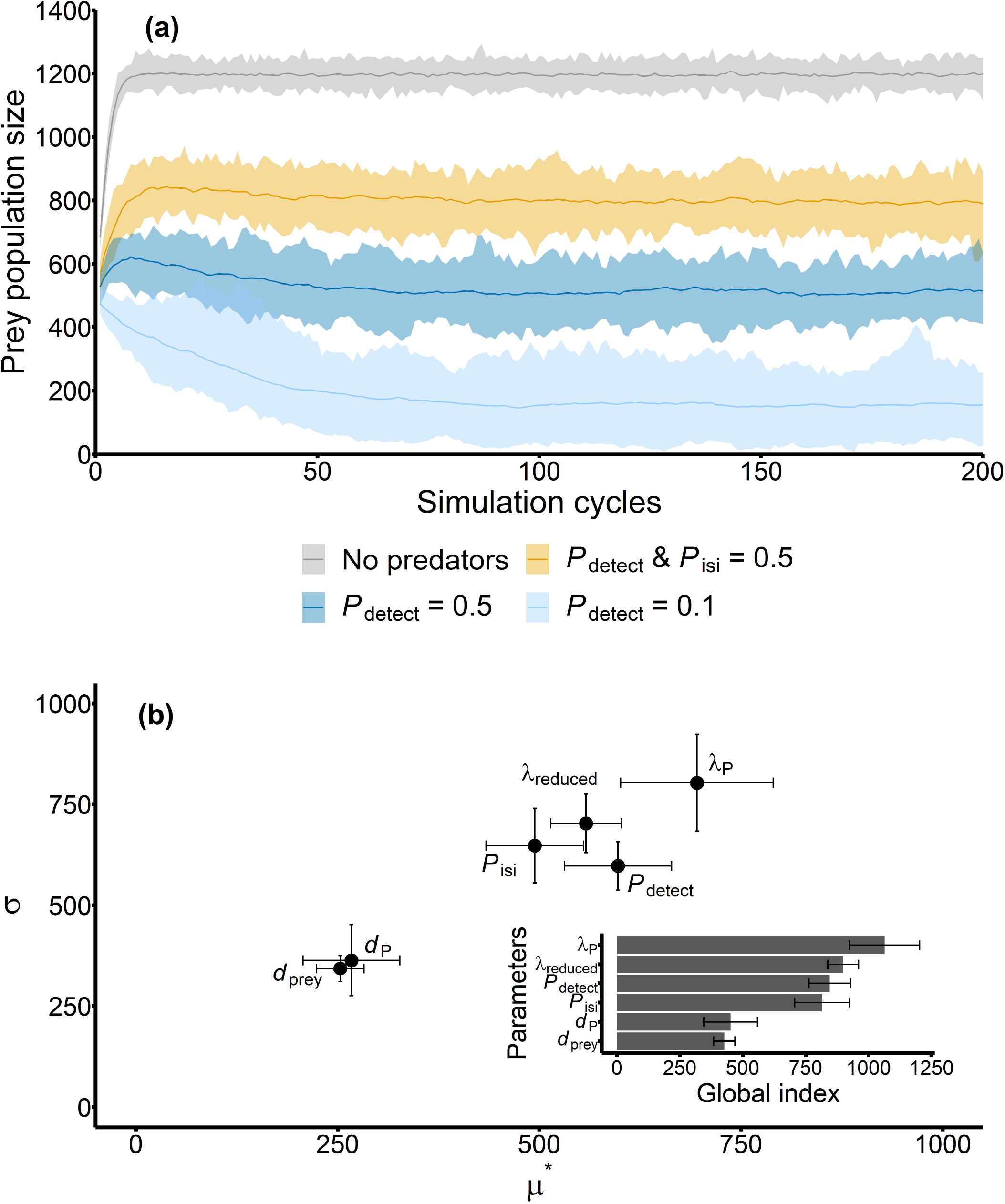
Effects of the *P*_detect_ and *P*_isi_ model parameters on the prey population. (a) Temporal fluctuations in prey abundance (means with range) without predators (grey) and under nominal parameter settings (with nominal *P*_detect_ and *P*_isi_ – orange, with nominal *P*_detect_ – dark blue, with minimal *P*_detect_ – light blue). When the predator detection probability was set to its minimal value, the prey population died out in a single iteration; in all other cases, the number of iterations was set to 50. (b) Results of the global sensitivity analysis (SA) depicting the impact of each model parameter on the mean (*x*-axis) and standard deviation (*y*-axis) of prey abundance; mean ± SD values for each parameter were calculated from five independent SA runs. Inset shows the model parameters ordered according to their overall influence on the model output

We characterised prey population sizes by calculating the mean, standard deviation, maximum and minimum values in four settings: in the absence of predators, with minimal *P*_detect_, with nominal *P*_detect_, and with nominal *P*_detect_ and *P*_isi_ parameter values, respectively. All other parameters were set to their initial values; for each model type, simulations were iterated 50 times. When the predator detection probability was set to its minimal value, the prey population died out in a single iteration; prey extinction was not observed in other settings.

We examined how predator abundance affected mortality rate due to predation in prey in the presence of minimal *P*_detect_, nominal *P*_detect_, and nominal *P*_detect_ and *P*_isi_ parameter values, respectively. All other parameters were set to their initial values. In each setting, simulation runs were iterated 50 times. If the prey population died out before the 200^th^ simulation cycle, the given run was omitted from the dataset (*n*=209; only in the ‘minimal predator detection’ setting).

#### Sensitivity analysis

We used Morris’s “OAT” elementary effects screening method (Morris 1991) with the extension introduced by Campolongo et al. (2007) as a global sensitivity analysis (SA) to rank the model parameters according to their impact on prey population size. We chose this SA because it produces results comparable to the more complex methods (Confalonieri et al. 2010) and is applicable to uncover the mechanisms and patterns produced by individual-based models (Imron et al. 2012; Beaudouin et al. 2015; ten Broeke et al. 2016). The mean of the absolute value of the elementary effect (*μ*^*^_*i*_) provides a measure for the overall influence of each input variable on the model output, whereas the standard deviation of the elementary effect (*σ_i_*) indicates possible non-linear effects or interactions among variables (Campolongo et al. 2007; Iooss and Lemaître 2015). We also ranked the model parameters using a global index (GI) (Ciric et al. 2012) calculated as:

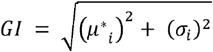

For the space-filling sampling strategy proposed by Campolongo et al. (2007), we generated *r*_2_ = 1000 Morris trajectories and then retained *r*_1_ = 50 with the highest ‘spread’ in the input space to calculate the elementary effect for each model parameter.

#### Parameter space exploration

We explored a specific part of the parameter space by visualising the combined effect of the parameters *P*_detect_, *P*_isi_, *λ*_P_ and *λ*_reduced_ on prey population size. Specifically, we investigated the effect of ISI use at low, intermediate and high levels of predator detection probabilities. In each scenario, predator avoidance behaviour had either no cost or incurred moderate fitness cost (i.e., decreased by one third compared to the maximum) and predation pressure was either low (0.025), intermediate (0.05) or high (0.075). In each setting, we used the complete range of parameter values for *P*_isi_ (Table 1). Simulations were iterated 30 times. In the low predator detection probability scenario coupled with high predation pressure, the prey population died out in the majority of simulation runs (*n*=581); these simulation outputs were omitted from the assembled dataset.

#### Network characterisation

We generated network data by running the model with *λ*_P_=0.075 (i.e., under a high level of predation pressure), while all other parameters were set to their nominal values. We compared emerging detection networks that were generated with the presence of ISI use (*P*_isi_=0.5) to those that were obtained when *P*_isi_=0. We calculated the number of components, component size, average ego network and average global efficiency as structural network properties for network characterisation. Simulations were repeated 50 times in each parameter setting. Additionally, we calculated the same characteristics for randomised detection networks as well (Farine 2017; Hobson et al. 2021). These were constructed from both type of the observed detection networks by randomly reshuffling the edges between nodes while also retaining the original degree distributions. Thus, randomized networks represented a hypothetical scenario where interactions are equally likely between any pair of nodes (Croft et al. 2011). While the main purpose here was to explore the global structure of the observed detection networks, the randomized networks helped us to assess whether ISI use could similarly affect the structural properties of networks that were based on this simplifying assumption. The number of components represents the number of connected parts in the detection networks (isolated nodes excluded). We computed component size as the number of components divided by the number of connected nodes; this measure denotes the average number of nodes embedded within components. We calculated the average size of ego networks as the mean number of reachable nodes within two steps in the components. To estimate transmissibility within components, we used the measure ‘global efficiency’ (Latora and Marchiori 2001; Pasquaretta et al. 2014; Romano et al. 2018). Global efficiency for a graph with *N* vertices is:

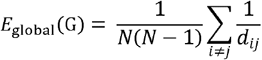

where *d*_ij_ is the shortest path length between nodes *i* and *j*. The value of this measure ranges from 0 to 1, and represents how fast information may spread from the source to the most peripheral network positions with the least number of connections (Romano et al. 2018). We computed global efficiency for the largest components in the networks. For the calculation of the above network properties, we used the ‘igraph’ and ‘brainGraph’ R packages (Csardi and Nepusz 2016; Watson 2020).

## Results

We found that nominal predation pressure coupled with minimal predator detection probability (*P*_detect_=0.1) led to small prey population size with high variation among runs compared to the null model when predators were absent and prey population existed at the carrying capacity of the environment (Fig. 3a, Table 2). Nominal predator detection probability (*P*detect=0.5) increased mean prey population size and stabilised the prey population at higher abundance values, while in the presence of nominal probability of ISI use in prey (*P*_detect_=0.5, *P*_isi_=0.5), prey population size increased further by approx. 53%. The sensitivity analysis also confirmed that *P*_isi_ was an influential model input in the constructed model (Fig. 3b). As expected, the parameters driving antipredator behaviour, i.e. the level of predation pressure, the probability of predator detection directly or via conspecifics, and the cost associated with performing antipredator behaviour, were all important and characterised by non-linear effects on prey abundance and/or strong interactions with other parameters. The parameters *d*_prey_ and *d*_P_ had considerably less influence on the dispersion of the model output, and were fixed to their nominal values in the subsequent analyses. The mechanism behind the effect of *P*_isi_ was that the presence of ISI use could decrease the per capita mortality due to predation across the whole range of the examined predation pressure regime and substantially mitigate the positive relationship between predation-related mortality rate and predator population size (Fig. 4, Fig. S1).

**Fig. 4.**
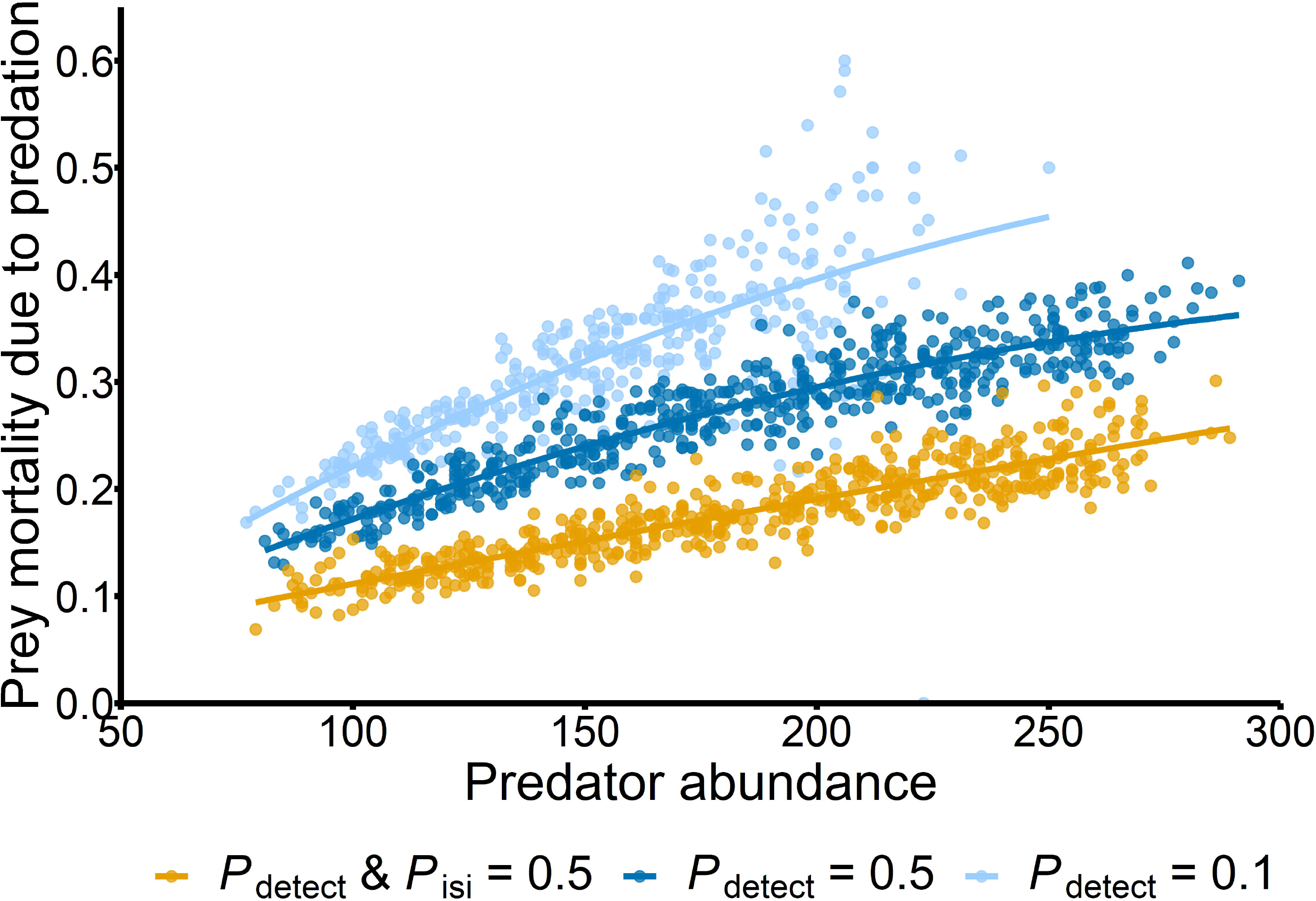
The relationship between per capita mortality due to predation and the number of predators using the same parameter settings as in Fig. 3a (but without the ‘No predators’ group). Trend lines were fitted using second-order polynomial approximation. Simulation results from incomplete runs (i.e., simulation cycles were less than 200) were omitted from the dataset (*n*=204; only in the ‘minimal predator detection’ model type)

**Table 2.**
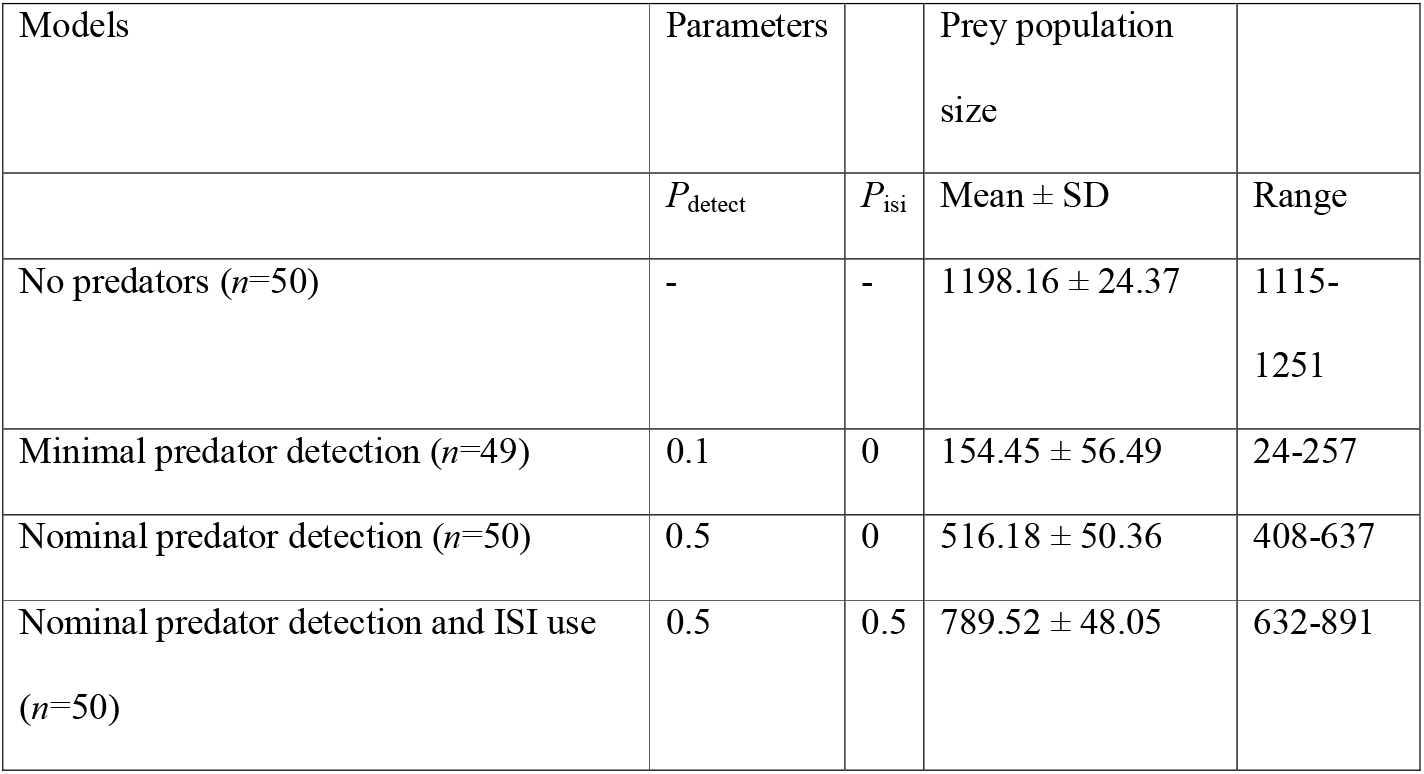
Descriptive statistics of the simulated prey populations calculated from 50 replicates at the 200^th^ simulation cycle (49 in the case of the second model type as prey population died out in a single iteration)

Consistent with expectations, *P*_isi_ affected prey number in all examined *P*_detect_ scenarios in interaction with the effect of cost and predator pressure (Fig. 5). This relationship was positive and nonlinear in most cases. When the predation pressure was low, *P*_isi_ positively influenced prey abundance to a limited extent, while the effect of the associated cost, especially at lower *P*_isi_ values, depended on the value of *P*_detect_. When the predation pressure was intermediate or high, *P*_isi_ exerted a more substantial influence on prey abundance and had the capacity to double the number of prey individuals irrespective of the presence or absence of associated cost (Table S1). Importantly, ISI use could counteract high predation pressure only when *P*_detect_ had a sufficient value (directly dependent on the degree of predation pressure), and did not compensate for low predator detection ability as indicated by the high prevalence of population extinctions in prey when high predation pressure was coupled with low predator detection ability. The presence of associated fitness cost in the high predation pressure settings greatly reduced the magnitude of the effect of ISI use on prey population size, but *P*_isi_ could still increase prey population size even at intermediate values if *P*_detect_>0.25.

**Fig. 5.**
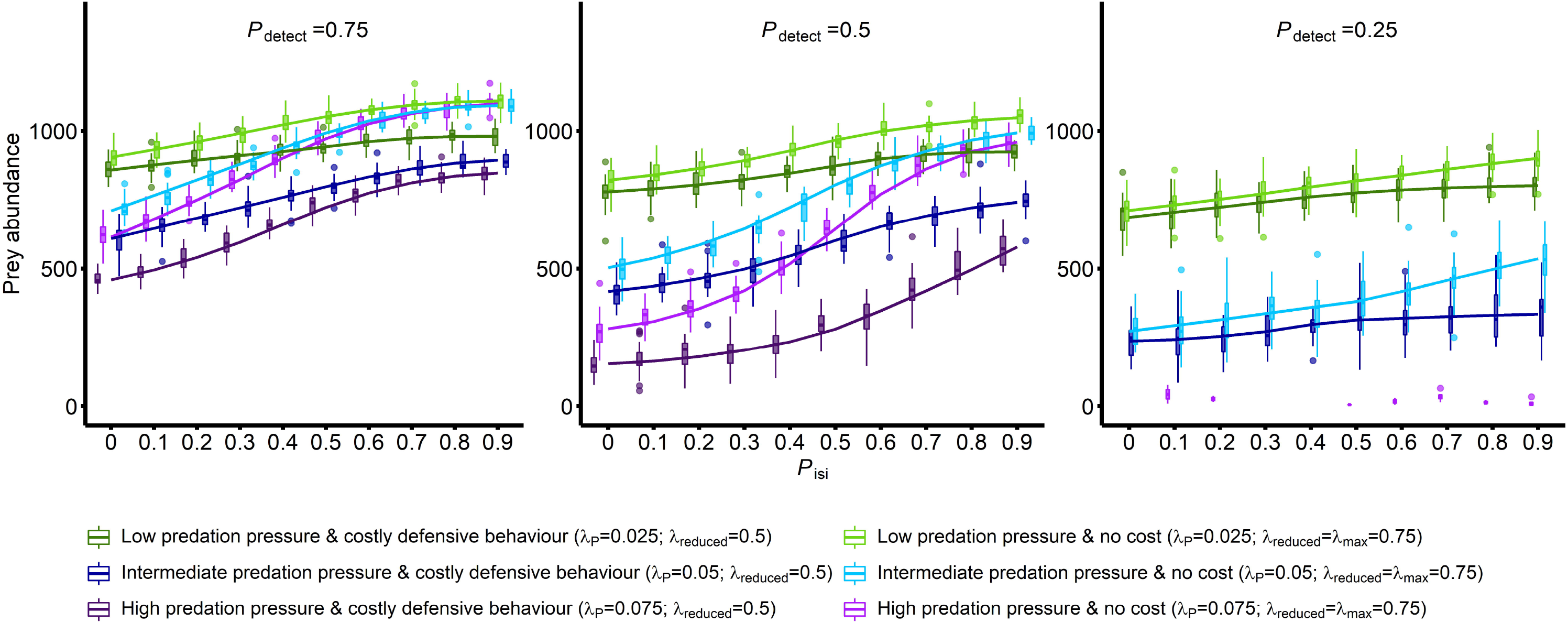
Interactive effects of the probability of ISI use (*P*_isi_), predation pressure (*λ*_P_) and the presence of fitness cost (associated with the defensive behaviour; *λ*_reduced_) on prey population size in three *P*_detect_ scenarios. The colour of the boxplots indicates the level of predation pressure (purple: high, blue: intermediate, green: low), while the colour tone is associated with the presence of cost (dark: costly defensive behaviour, light: no cost). Trend lines were fitted using the ‘LOESS’ regression method for smoothing with the default value of span (0.75); presented only for illustration purposes. Simulation results from incomplete runs were omitted from the dataset (*n*=581; only in the ‘*P*_detect_=0.25’ setting)

The observed detection networks were characterized by high numbers of components that consisted of few connected individuals and small ego networks (Fig. 6, Table S2). The number of connected individuals more than tripled when social information could spread among individuals, while the number of isolates did not change with ISI use. Twice as much components were found in the observed detection networks in the presence of ISI use compared to the setting when it was absent; this effect, however, was not detectable in the randomized counterparts. Mean component size was unaffected by the presence of ISI use in the observed networks, but increased substantially in the randomized ones. Ego network sizes were similarly influenced by ISI use in both network types. Global efficiency within the largest components was high in the absence of ISI use in both observed and randomized networks; however, it was also high in the presence of ISI use in the observed detection networks, indicating efficient information transmission among individuals whenever connected prey was able to detect nearby predators. These attributes of functioning detection networks were unlikely to be the direct consequence of higher prey population size in the presence of ISI use, because the corresponding randomized networks did not show the same degree of structural changes compared to the *P*_isi_=0 setting.

**Fig. 6.**
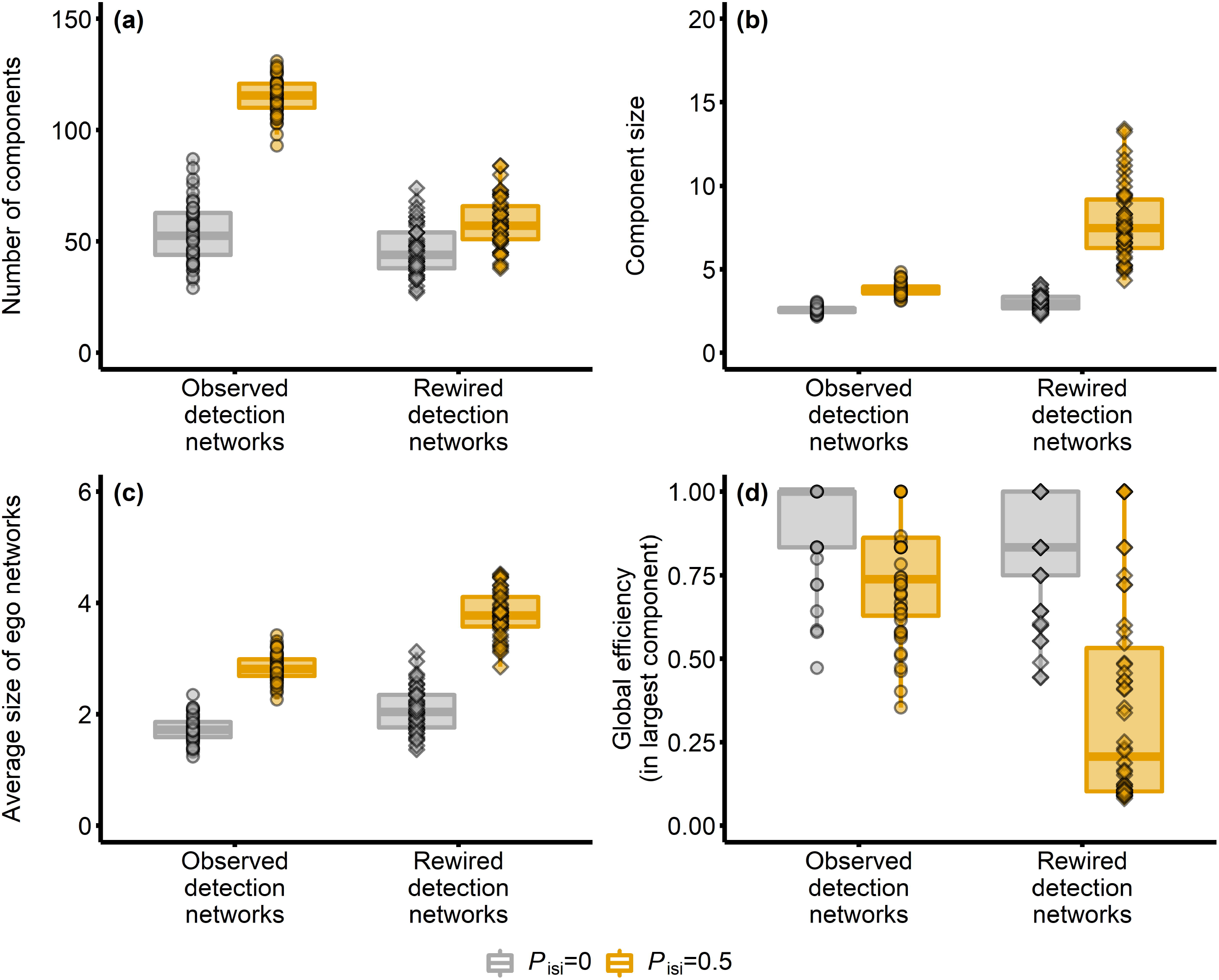
Four structural network properties (a: number of components, b: component size, c: average size of ego networks, d: average global network efficiency) calculated for the observed detection networks (circles) and corresponding randomized networks (diamonds). Predation pressure was set to ‘high’ (i.e., *λ*_P_=0.075). Boxplots show the median and interquartile range, whiskers denote values within 1.5-fold of the interquartile range, and dots are individual values. The colour of the boxplots indicates the absence (gray; *P*_isi_=0) or presence of ISI use (orange; *P*_isi_=0.5)

## Discussion

Social information use has been assumed both to increase individual fitness and to affect population- and community-level processes (Dall et al. 2005; Gil et al. 2018). We expected that such effects could emerge in randomly moving non-grouping prey if behavioural contagion can occur through detection networks, i.e., a dynamic system of temporary observation-based connections between conspecifics. Correlated random walk has successfully been used to describe non-oriented movement trajectory in a number of non-grouping animals, e.g. insects (e.g., Kareiva and Shigesada 1983; McCulloch and Cain 1989; Byers 2001), zooplankton (e.g., Komin et al. 2004; Uttieri et al. 2004, 2008), echinoderms (e.g., Lohmann et al. 2016) or mammals (e.g., Johnson et al. 2008). We found that irrespective of the apparent stochasticity in our model, the sharing of adaptive antipredator behaviour could contribute to population stability and persistence in prey by mitigating predation-related per capita mortality and raising equilibrium population sizes. We also showed that temporary detection networks had structural properties that allowed the efficient spread of adaptive antipredator behaviour among prey under high predation pressure. In group-living animals, information spreads via social connections among individuals and social network positions strongly interact with individual spatial behaviour (Firth and Sheldon 2016; Spiegel et al. 2016; Webber and Vander Wal 2018; Albery et al. 2021), thus movement characteristics and space use are shaping information transmission by affecting social connections. Our findings indicate that non-grouping animals, by being embedded in detection networks based on their perception attributes and spatial locations, can benefit from similar information transmission processes as well. As inadvertent social information use in non-grouping animals is largely understudied (see in Tóth et al. 2020), in the following paragraphs we contrasted our simulation results on non-grouping prey with previous (empirical) findings on group-living species in several instances.

Our results corroborate with previous studies on group-living organisms indicating that social information may act as a stabilizing mechanism in systems where predators can exert high pressure on prey populations (Gil et al. 2017, 2018, 2019). While in those models social information directly reduced (following a specific function) the per capita mortality (e.g., Gil et al. 2018), the presented work offers a more mechanistic understanding of how inadvertent social information could propagate through a population of randomly moving individuals. Our findings indicate that predator detection ability had to reach a sufficient level, strongly dependent on the actual level of predation pressure, for ISI use to facilitate prey population persistence. Notably, when this condition was met, ISI use exerted a detectable positive influence on prey population size by relaxing predation pressure even at low probabilities and even if the adaptive antipredator behaviour incurred a fitness cost. Although the depth of our understanding of the detected non-linear relationships and potential thresholds is limited by their coarse-grained variation in these parameters examined here, simulations nonetheless prove that in a substantial part of the parameter space social information use can be expected to raise non-grouping prey population size and facilitate its persistence. These findings may have crucial implications in many theoretical and applied ecological contexts, ranging from the invasive dynamics of predator-prey systems to the efficiency of biological control practices. For instance, the recognition of novel predators by naïve prey has been associated with social information use via different perception modalities in group-living fish (Ferrari et al. 2005; Manassa et al. 2013), and similar utilization of social cues in social birds has been shown to facilitate the spread of novel aposematic prey (Thorogood et al. 2017; Hämäläinen et al. 2021). Such social information-mediated interactions between prey and predators might be more prevalent in natural ecosystems that include non-grouping species as well, contributing to deviations from the predictions of theoretical models in the dynamics of trophic interactions (Polis et al. 2000). When natural enemies are used as biological control agents for pest management, diffusion of antipredator responses among prey may substantially reduce predation rates rendering these practices less effective and profitable. Besides, it may also mitigate the expected positive impact of the non-consumptive effects of predators (NCEs; Preisser et al. 2007; Sih et al. 2010) such as decreased crop damage due to reduced feeding rate in pests (Beleznai et al. 2017; Tholt et al. 2018). This inflation of NCEs due to information spread can generate discrepancies in the findings of large-scale field studies and laboratory experiments (see in Weissburg et al. 2014), and should be taken into consideration in investigations that aim to evaluate how NCEs may trigger trophic cascades in different ecosystems (Herman and Landis 2017; Haggerty et al. 2018; Pessarrodona et al. 2019).

Detection networks had distinct structural characteristics when prey experienced high predation pressure and exploited social cues to avoid predators. These networks typically consisted of many components with few connected individuals and small average ego networks, and within these small components, social information could spread with relatively high efficiency. The key to understanding the differences in structural properties of detection networks in the presence and absence of ISI use lies in identifying the process that generates more and smaller components. One plausible explanation is that prey distribution in the simulated landscape could remain more homogeneous due to a decreased susceptibility to predation in the vicinity of predators as the diffusion of social information greatly enhances the probability of predator detection even among a few nearby individuals. While high network efficiency has previously been identified in small animal groups, cognitive abilities and strong social affiliations have usually been involved in explaining this emergent property (Waters and Fewell 2012; Pasquaretta et al. 2014). Our findings indicate that incidental connections among non-grouping animals may generate networks that have similar favourable attributes. In addition to differences in the sizes of connected components, there may be other key differences in how information spreads through detection or sensory networks among group-living (Strandburg-Peshkin et al. 2013; Rosenthal et al. 2015; Davidson et al. 2021) and non-grouping individuals, however. First, behavioural contagion can be complex, and the number of non-alarmed individuals within the detection range influences the likelihood of adopting a specific behaviour (Firth 2020). Previous works on social species have provided mounting evidence for such complex contagion (Hoppitt and Laland 2013; Grüter and Leadbeater 2014; Kendal et al. 2018). Second, imperfect copying might decrease the intensity of behavioural responses with each transmission step, and under a given threshold intensity, social cues exert no response from nearby observers. In this case, individuals’ ability to convey information about predation hazards is related to the extent of behavioural change compared to a baseline level (Chivers and Ferrari 2014). Third, phenotypic heterogeneity among individuals may influence information diffusion if individual traits (e.g., related to hunger, age or developmental stage) or functional traits that transcend species (e.g., similarity in body size that may lead to shared predators) affects the individual capacity to produce social information (Farine et al. 2015).

To describe how ISI use may affect population dynamics in non-grouping prey, we constructed a tentative model with naturalistic predator-to-prey ratios (1:1.03 [when predator detection probabilities was set to minimal]–1:4.23 [with nominal predator detection and ISI use probabilities]; see in Donald and Anderson 2003). Previous observations indicate that predator detection probability, which has been found to play a crucial role in the emergence of social information-mediated effects in our study, can have a value within the upper half of the range examined here (i.e., >0.5) under relevant conditions (e.g., Tisdale and Fernández-Juricic 2009; Manzur et al. 2018). However, being strongly dependent on the neuronal pathways underlying detection mode and the processing capacity of the brain (Clark and Dukas 2003; Pereira and Moita 2016), it can differ significantly between species and even within the same species as it may also depend on the forager’s state of energy reserves (Clark and Mangel 2000). Therefore, to construct a more realistic model, both species-specific and context-specific information (e.g., movement distances, detection ranges and reproduction rates) for existing predator-prey relationships need to be incorporated, which can be done only at the expense of generality. Model precision may be further enhanced by incorporating additional variables including the functional response of specific predator species (Dunn and Hovel 2020), different non-consumptive effects (other than reduced feeding rate) (Peckarsky et al. 2008), a measure of social cue reliability (Dunlap et al. 2016), social information use in predators (Hämäläinen et al. 2021), or landscape heterogeneity that could alter the space use of individuals (Albery et al. 2021). The effects of different transmission modes can also be tested, for instance, by weighting the probability of information diffusion among conspecifics by the proportion of alarmed and non-alarmed individuals within the detection zone or incorporating heterogeneity among individuals in attributes that affect their propensity to act as social cue producers. Our work, thus, represents a general modelling approach that could be applied to predator-prey systems in which populations are demographically decoupled and non-grouping prey may mitigate predation hazards through the exploitation of incidentally produced social information.

## Acknowledgments

We are indebted to Béla Keresztfalvi for providing access to the institutional IT resources, with which we were able to reduce the computational time of the performed simulations radically. We are also thankful to the reviewers for their constructive comments.

## Statements and Declarations

### Ethics approval

Not applicable.

### Data availability

Data files supporting the results and R script for model construction and simulated data are archived and available at Figshare (https://figshare.com/s/34fc714342dab9123193). Upon reasonable requests, R codes for the model functions are also available from the corresponding author.

### Authors’ contributions

ZT and CG conceived and designed the study. ZT constructed the model, performed the simulations, analysed the model output, wrote the initial manuscript, and revised and edited the subsequent versions. CG contributed substantially to the text and revisions.

