## Supplementary material for "Social information-mediated population dynamics in non-grouping prey": Fig. S1

submitted to Behavioral Ecology and Sociobiology

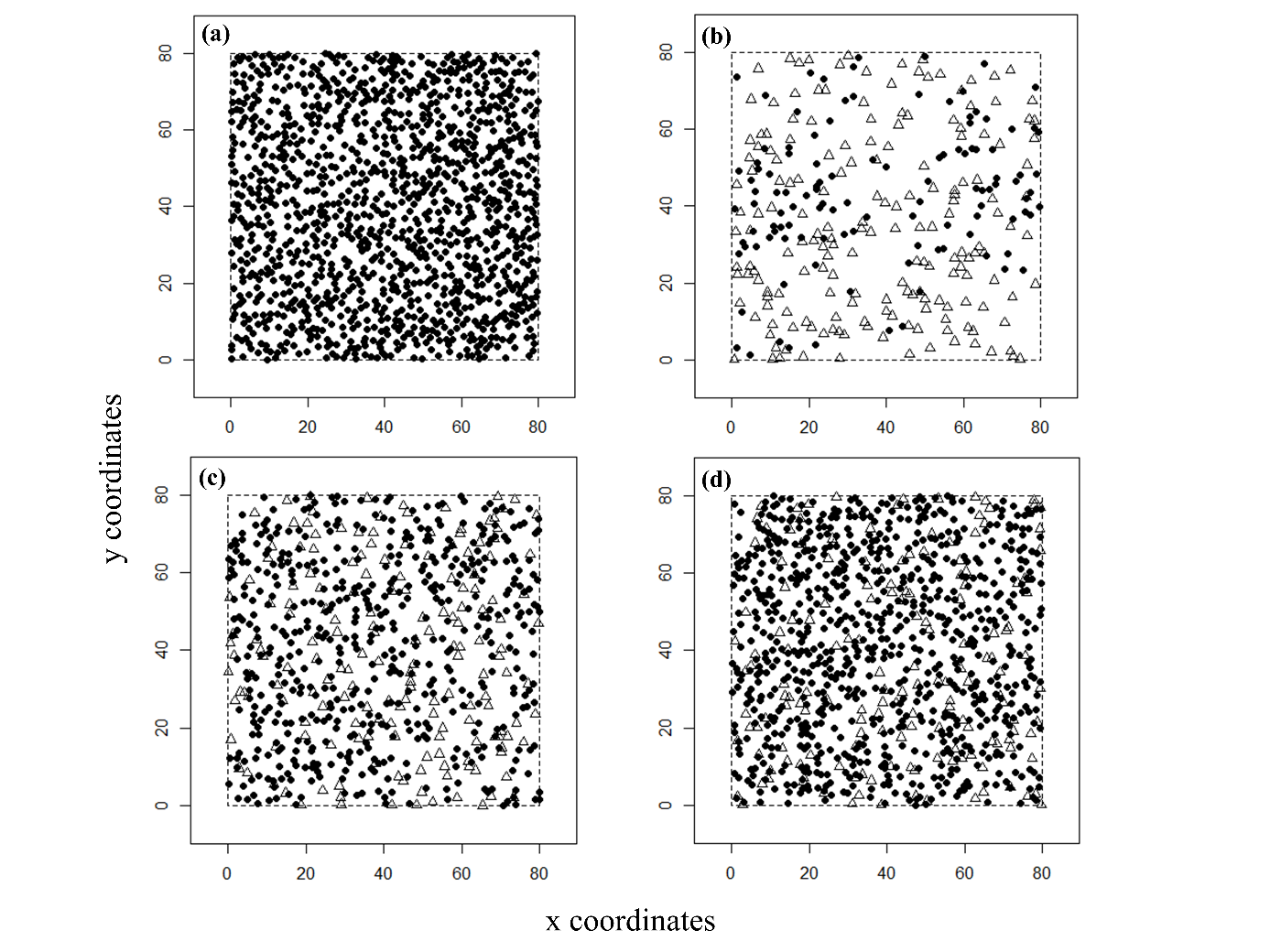

Figure S1. Examples of individual locations (filled circles: prey, empty triangles: predators) on the landscape at the 200th simulation cycle for each of the four scenarios depicted in Figure 3a (a: in the absence of predators, b: with minimal *P*_detect_, c: with nominal *P*_detect_, d: with nominal *P*_detect_ and *P*_isi_ parameter values).

Table S1. Prey population sizes (mean and SD) in different combinations of the four model parameters *P*_detect_ (predator detection probability), *P*_isi_ (probability of ISI use), *λ*_P_ (predation pressure) and *λ*_reduced_ (cost of antipredator behaviour).

|  |  |  | *P*_isi_=0 |  | *P*_isi_=0.5 |  | *P*_isi_=0.9 |  |
| --- | --- | --- | --- | --- | --- | --- | --- | --- |
|  |  |  | Mean | SD | Mean | SD | Mean | SD |
| Low predation pressure (*λ*_P_=0.025) | Costly behaviour (*λ*_reduced_=0.5) | *P*_detect_ =0.25 | 690.03 | 63.49 | 781.53 | 49.85 | 801.3 | 48.77 |
|  |  | *P*_detect_ =0.5 | 778.67 | 51.06 | 871.93 | 48.93 | 922.43 | 30.77 |
|  |  | *P*_detect_ =0.75 | 860.1 | 35.54 | 938.83 | 29 | 979.4 | 34.84 |
|  | No cost (*λ*_reduced_=0.75) | *P*_detect_ =0.25 | 710.17 | 58.01 | 814.8 | 60.41 | 898.77 | 49.12 |
|  |  | *P*_detect_ =0.5 | 819.6 | 54.72 | 960.83 | 33.38 | 1050.5 | 35.81 |
|  |  | *P*_detect_ =0.75 | 905.93 | 34.94 | 1050.4 | 29.39 | 1107.1 | 38.83 |
| Intermediate predation pressure (*λ*_P_=0.05) | Costly behaviour (*λ*_reduced_=0.5) | *P*_detect_ =0.25 | 237.37 | 58.88 | 327.7 | 87.96 | 334.07 | 81.01 |
|  |  | *P*_detect_ =0.5 | 411.6 | 53.83 | 597.93 | 51.09 | 743.3 | 43.93 |
|  |  | *P*_detect_ =0.75 | 607.23 | 54.24 | 793.1 | 33.26 | 891.43 | 27.07 |
|  | No cost (*λ*_reduced_=0.75) | *P*_detect_ =0.25 | 279 | 60.54 | 390.13 | 87.59 | 526.9 | 79.99 |
|  |  | *P*_detect_ =0.5 | 500.5 | 54.22 | 804.67 | 46.11 | 995.93 | 31.04 |
|  |  | *P*_detect_ =0.75 | 712.5 | 40.3 | 992.3 | 24.81 | 1090.73 | 31.41 |
| High predation pressure (*λ*_P_=0.075) | Costly behaviour (*λ*_reduced_=0.5) | *P*_detect_ =0.25 | - | - | - | - | - | - |
|  |  | *P*_detect_ =0.5 | 150 | 44.23 | 299.3 | 52.65 | 570.83 | 60.84 |
|  |  | *P*_detect_ =0.75 | 462.87 | 27.96 | 729.53 | 40.47 | 848.57 | 32.62 |
|  | No cost (*λ*_reduced_=0.75) | *P*_detect_ =0.25 | - | - | 5.5 | 0.71 | 12.25 | 14.59 |
|  |  | *P*_detect_ =0.5 | 271.17 | 54.99 | 645.93 | 40.59 | 956.8 | 41.7 |
|  |  | *P*_detect_ =0.75 | 618.6 | 45.69 | 978.1 | 32.68 | 1103.2 | 25.48 |

Table S2. Descriptive statistics of the examined detection network properties calculated under a high level of predation pressure (*λ*_P_=0.075) in the absence (*P*_isi_=0) or presence of ISI use (*P*_isi_=0.5). B: both network types, O: observed networks, R: randomized networks.

| Structural characteristics | Network type | *P*_isi_=0 |  |  |  | *P*_isi_=0.5 |  |  |  |
| --- | --- | --- | --- | --- | --- | --- | --- | --- | --- |
|  |  | Mean | SD | Min | Max | Mean | SD | Min | Max |
| Connected individuals | B | 138.14 | 41.3 | 65 | 247 | 433.96 | 49.15 | 292 | 536 |
| Isolates | B | 133.38 | 10.55 | 107 | 161 | 119.66 | 10.25 | 101 | 145 |
| Number of components | O | 53.64 | 13.37 | 29 | 87 | 115.1 | 8.09 | 93 | 131 |
|  | R | 45.78 | 10.78 | 27 | 74 | 58.26 | 11.98 | 36 | 87 |
| Mean component size | O | 2.55 | 0.2 | 2.16 | 3.03 | 3.77 | 0.37 | 3.10 | 4.83 |
|  | R | 3 | 0.47 | 2.26 | 4.19 | 7.88 | 2.35 | 4.28 | 13.29 |
| Mean ego network size | O | 1.73 | 0.24 | 1.24 | 2.35 | 2.84 | 0.24 | 2.26 | 3.42 |
|  | R | 2.08 | 0.4 | 1.37 | 3.13 | 3.79 | 0.41 | 2.84 | 4.52 |
| SD of ego network size | O | 0.94 | 0.23 | 0.52 | 1.6 | 1.60 | 0.15 | 1.18 | 1.9 |
|  | R | 1.2 | 0.35 | 0.58 | 2.25 | 2.24 | 0.25 | 1.74 | 2.84 |
| Global efficiency within largest component | O | 0.91 | 0.13 | 0.47 | 1 | 0.75 | 0.18 | 0.35 | 1 |
|  | R | 0.81 | 0.19 | 0.38 | 1 | 0.36 | 0.32 | 0.08 | 1 |
